# Modeling the Role of Hypoxia in Glioblastoma Growth and Recurrence Patterns

**DOI:** 10.1101/691238

**Authors:** Lee Curtin, Andrea Hawkins-Daarud, Alyx B. Porter, Markus R. Owen, Kristoffer G. van der Zee, Kristin R. Swanson

## Abstract

A typical feature of glioblastoma (GBM) growth is local recurrence after surgery. However, some GBMs recur distally. It has been noted that GBM patients with perioperative ischemia are more likely to have distal recurrence and that GBM cells migrate faster under hypoxic conditions. We apply the Proliferation Invasion Hypoxia Necrosis Angiogenesis (PIHNA) model to examine the effect of faster hypoxic cell migration on simulated GBM growth. Results suggest that a highly migratory hypoxic cell population drives the growth of the whole tumor and leads to distant recurrence, as do higher normoxic tumor cell migration and low cellular proliferation rates.

## I. Introduction

Glioblastoma (GBM) is the most aggressive primary brain tumor with a median survival of 15 months [1]. GBM tumors are very invasive, migrating in with healthy tissue in low densities at their leading edge beyond the resolution of current magnetic resonance imaging (MRI). Typically, the first stage of treatment is surgery. However, GBM cells remain and repopulate, often from the leading edge – which is known as a local recurrence. It has been noted that tumors of patients who suffer from vascular ischemia in the brain following surgery recur far from the leading edge or more diffusely – which are known as distant and diffuse recurrences, respectively [2]. This has been attributed to the hypoxic conditions created by the local ischemia, but the mechanism by which this occurs is not fully understood [2]. There is also evidence that tumor cells exhibiting signs of hypoxia migrate at a faster rate [3]. To investigate how faster hypoxic migration rates affect the growth dynamics of simulated GBMs and their recurrence patterns, we utilize the Proliferation Invasion Hypoxia Necrosis Angiogenesis (PIHNA) mathematical model of GBM growth. We apply the PIHNA model, which incorporates the angiogenic interactions between the tumor and vasculature on a tissue scale [4], to a case of perioperative ischemia and look for mechanisms that promote simulated distant recurrence.

## II. Methods

### A. The PIHNA Model

The PIHNA model simulates five species: normoxic cells, hypoxic cells, necrotic cells, vascular endothelial cells and angiogenic factors. Normoxic cells migrate with rate *D*_*c*_, proliferate with rate *ρ*, create angiogenic factors, and become hypoxic under low-vasculature conditions. Hypoxic cells migrate with rate *D*_*h*_ and become necrotic under low-vasculature conditions, they create angiogenic factors at a larger rate than normoxic cells to encourage the creation of vasculature at their location. Necrotic cells accumulate and encourage other local cell types to become necrotic. These dynamics produce a spread of normoxic cells leading the tumor growth, followed by hypoxic and necrotic cells towards the core of the tumor, which agrees with clinical observations [4]. Vascular cell density begins at a homeostatic level, increases near the hypoxic cells (mediated through their release of angiogenic factors) and decreases towards the necrotic core of the tumor.

### B. Wave-speed Analysis of the PIHNA Model

We carried out a wave-speed analysis on the spherically symmetric PIHNA model, analogous to that of Fisher’s Equation [5]; this gives the model parameters that influence the growth rate of the whole tumor.

We run spherically symmetric PIHNA simulations to verify the predicted minimum wave-speed, with different rates of *D*_*c*_ and *ρ*. We also vary the relative migration rates between normoxic and hypoxic cell populations, *D*_*h*_/*D*_*c*_, to investigate the effect of a faster-migrating hypoxic cell population on the tumor wave-speed.

### C. Simulating Resection and Perioperative Ischemia with the PIHNA Model

We use the geometry from the BrainWeb Database [6] that delineates different brain regions to run PIHNA simulations. We allow for growth in white and gray matter regions, but not the skull or cerebrospinal fluid regions.

Following previous work, we assume that a total cell density of 80% is the threshold required for a tumor to be visible on a T1-weighted magnetic resonance image with contrast (T1Gd MRI) and 16% to be visible on T2-weighted imaging (T2 MRI) [7].

We simulate a lobectomy that removes the tumor bulk on simulated T1Gd MRI and a transient ischemic reduction to a region next to the resection cavity. We then allow the tumor to continue growing according to the model equations and classify tumor recurrence location as the first appearance on simulated T1Gd MRI of a tumor mass above a threshold size. We run these simulations for varying values of *D*_*c*_, *ρ*, and *D*_*h*_*/D*_*c*_. Figure 1 shows two simulations, one with and one without ischemia, where the resection and ischemic geometries can be seen.

**Figure 1.**
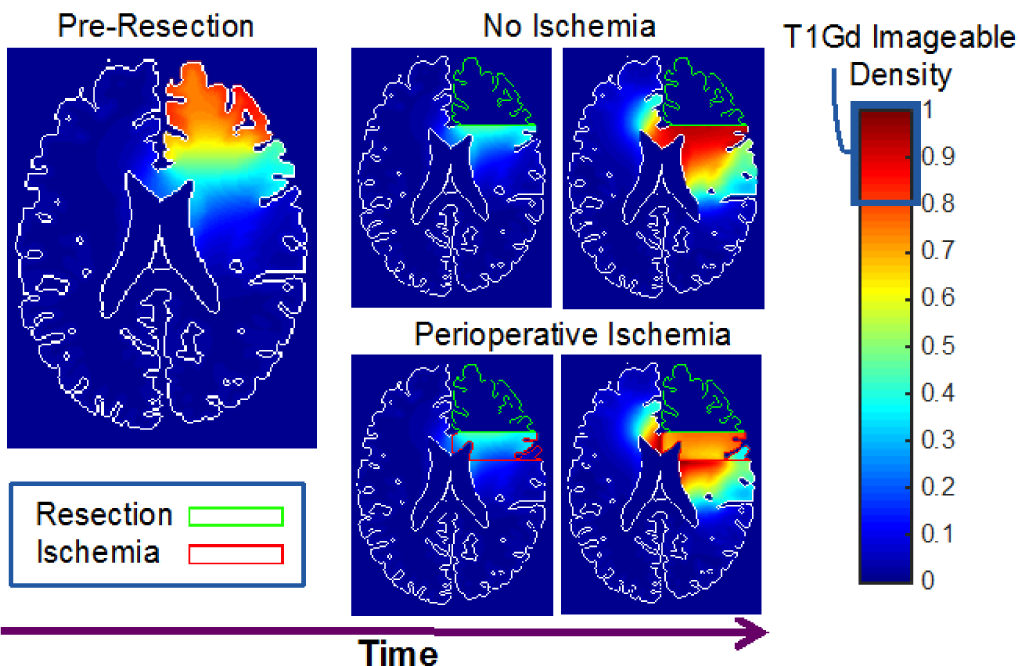
Total cell density for two example simulations, one with and one without perioperative ischemia. We see that the ischemia locally reduces the cell density to below the assumed T1Gd imageable density threshold, such that the tumor recurs as a distant recurrence close to the ventricles on simulated T1Gd MRI in the ischemia case, but locally recurs in the case without ischemia.

## III. Results

We find a minimum wave-speed of 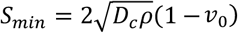, where *v*_0_ is the initial vasculature in the healthy brain. We see that this minimum wave-speed is attained in simulations if *D*_*c*_ is equal to or faster than the hypoxic counterpart, *D*_*h*_. In the case of large enough *D*_*h*_*/D*_*c*_, PIHNA simulations give a faster wave-speed, see Figure 2, suggesting that the faster-migrating hypoxic cell population can drive the outward growth of the tumor as a whole.

**Figure 2.**
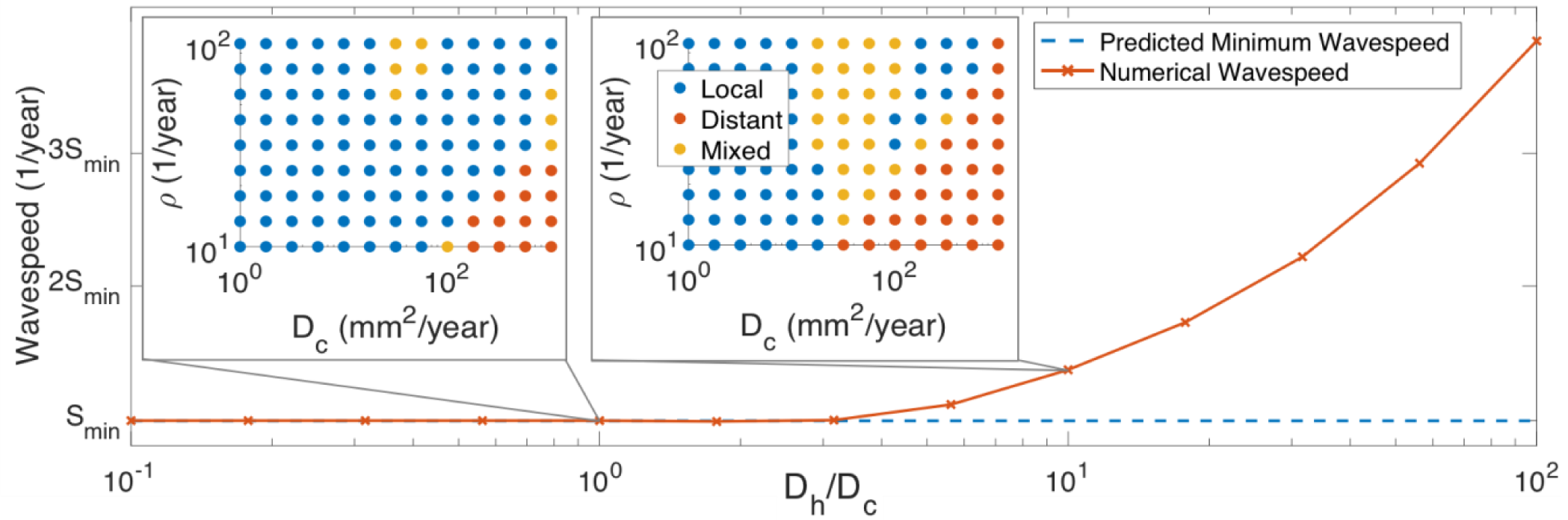
Simulations for higher *D*_*h*_*/D*_*c*_ lead to larger numerical tumour wave-speeds in 1D PIHNA simulations. Simultaneously, we see more distantly recurring tumours for these higher ratios, suggesting that the hypoxic cell diffusion drives the outward growth of the tumour and promotes distant recurrence.

The ischemia analysis suggests that high *D*_*c*_ rates, low *ρ* rates and high *D*_*h*_*/D*_*c*_ values all promote distant recurrence, the latter of which also increases the tumor wavespeed, see Figure 2. We see that the ischemic region reduces the local total cell density, such that the tumour recurs on simulated T1Gd MRI outside of this region, see Figure 1.

## IV. Discussion

We found the minimum analytical wave-speed for the PIHNA model, that only depends on the characteristics of the normoxic cells and the background vasculature in the brain. However, hypoxic cells that migrate at a faster rate than normoxic cells can increase the tumor wave-speed past the minimum and drive the outward tumor growth. We see that simulated tumors generally recur distantly if they have a high enough migration rate and low enough proliferation rate, and this is also promoted by hypoxic cells migrating faster than normoxic cells.

This analysis expands on the clinical observation that ischemia promotes distant recurrence by suggesting intrinsic tumor kinetics also play a role. The PIHNA model suggests that cells exhibiting a hypoxic phenotype of faster diffusion could be a target for therapies; if the hypoxic cell diffusion rate was inhibited, it could limit the outward growth rate of the tumor overall and lead to less observed distantly recurring tumors.

## References

[1] R. Stupp et al., “Effects of radiotherapy with concomitant and adjuvant temozolomide versus radiotherapy alone on survival in glioblastoma in a randomised phase III study: 5-year analysis of the EORTC-NCIC trial,” Lancet Oncol., vol. 10, no. 5, pp. 459–466, 2009.

[2] A.-L. Thiepold et al., “Perioperative cerebral ischemia promote infiltrative recurrence in glioblastoma,” Oncotarget, vol. 6, no. 16, p. 14537, 2015.

[3] O. Keunen et al., “Anti-VEGF treatment reduces blood supply and increases tumor cell invasion in glioblastoma,” Proc. Natl. Acad. Sci., vol. 108, no. 9, pp. 3749–3754, 2011.

[4] K. R. Swanson, R. C. Rockne, J. Claridge, M. A. Chaplain, E. C. Alvord, and A. R. Anderson, “Quantifying the role of angiogenesis in malignant progression of gliomas: in silico modeling integrates imaging and histology,” Cancer Res., vol. 71, no. 24, pp. 7366–7375, 2011.

[5] R. A. Fisher, “The Wave of Advance of Advantageous Genes,” Ann. Eugen., vol. 7, no. 4, pp. 355–369, 1937.

[6] D. L. Collins et al., “Design and construction of a realistic digital brain phantom,” IEEE Trans. Med. Imaging, vol. 17, no. 3, pp. 463–468, Jun. 1998.

[7] K. R. Swanson, C. Bridge, J. D. Murray, and E. C. Alvord Jr, “Virtual and real brain tumors: using mathematical modeling to quantify glioma growth and invasion,” J Neurol Sci, vol. 216, no. 1, pp. 1–10, Dec. 2003.

